# Ongoing emergence of M1_UK_ lineage among invasive group A streptococcus isolates in 2020 and use of allele-specific PCR

**DOI:** 10.1101/2022.12.18.520871

**Authors:** Xiangyun Zhi, Ho Kwong Li, Hanqi Li, Zuzanna Loboda, Samson Charles, Ana Vieira, Kristin Huse, Elita Jauneikaite, Juliana Coelho, Theresa Lamagni, Shiranee Sriskandan

## Abstract

**Background:** An increasing burden of invasive group A streptococcal infections is reported in multiple countries, notably England, where scarlet fever cases are also abundant. In England, increased scarlet fever and invasive infections have been associated with emergence of a sublineage of *emm*1 *Streptococcus pyogenes* that expresses increased SpeA scarlet fever erythrogenic toxin. Wider surveillance for toxigenic *Streptococcus pyogenes* lineage M1_UK_ is much needed however, to date, lineage assignment has required genome sequencing limiting surveillance to those centres with access to such facilities.

**Methods:** To circumvent the requirement for genome sequencing, an allele-specific PCR was developed to distinguish M1_UK_ from other *emm*1 strains. Additional PCR assays were developed to distinguish M1_UK_ from two intermediate lineages that were detected previously. The assay was evaluated using DNA from genome-sequenced upper respiratory tract *emm*1 *S. pyogenes* strains and a further set of 16 genome-sequenced invasive *S. pyogenes* isolates that included the two intermediate lineages. The assay was then applied to DNA from all 305 invasive *emm*1 isolates that had been submitted to the reference laboratory in the one pear period Jan 1-Dec 31 2020, in order to assign lineage.

**Results:** The allele specific PCR was 100% accurate when compared with genome sequencing, correctly identifying M1_UK_, two intermediate sublineages, and other *emm*1 strains. The assay demonstrated the M1_UK_ lineage to be dominant among *emm*1 invasive isolates in England, representing 278/305 (91%) of invasive *emm*1 isolates by end of 2020.

**Implications:** *Emm*1 *S. pyogenes* have a prominent role in invasive infections; any *emm*1 lineage that demonstrates enhanced fitness within the population is of public health concern. The allele specific PCR provides a readily available method to subtype *emm*1 isolates and does not require access to complex sequencing facilities. The data confirm that the M1_UK_ lineage has persisted and further expanded in England underlining the importance of wider global surveillance.

---

Upsurges in invasive group A streptococcal infection have been widely reported in England and elsewhere (1), emphasising a need for greater understanding of the relation between circulating *Streptococcus pyogenes* that cause pharyngitis and scarlet fever, and cases of invasive disease. While many factors such as exposure history, comorbidity, viral co-infection, genetic susceptibility may intersect to increase susceptibility to infection, strain-specific virulence may be important.

In England, where both scarlet fever and invasive *S. pyogenes* infections are notifiable, pronounced upsurges in scarlet fever have been recorded for the last 8 years albeit subsiding during the COVID-19 pandemic period (2, 3). During the 2015/2016 season, a notable increase in invasive infections was observed that had not been evident previously (4). Both scarlet fever and invasive infections were associated with the emergence of a new sublineage of *emm*1 *S. pyogenes* (M1_UK_) (4) that appeared to be outcompeting the highly successful contemporary epidemic *emm*1 clone (M1_global_) that had emerged and spread globally since the 1980’s (5, 6). Despite an unchanged phage repertoire, M1_UK_ strains were found to produce more superantigenic scarlet fever toxin SpeA (streptococcal pyrogenic exotoxin A) than contemporary M1_global_ *S. pyogenes* strains (3).

*Emm*1 *S. pyogenes* strains are known to be highly virulent (5) and disproportionately associated with invasive infections; any increase in the prevalence of *emm*1 strains in pharyngitis and scarlet fever is therefore of public health concern. Knowledge of the distribution of M1_UK_ is largely limited to those countries undertaking and reporting genome sequencing (Figure 1). M1_UK_ has been identified in other European countries, from a single isolate in Denmark (4) to more dominant status in the Netherlands (7). The lineage has also been reported in North America; the Public Health Agency of Canada reported 17/178 (10%) of *emm*1 isolates from 2016 to be M1_UK_ and broadly distributed across Canada (8). This contrasts with a reported M1_UK_ frequency of just 0 −2.8% of *emm*1 isolates in the USA, based on data from the Active Bacterial Core surveillance (ABCs) system of the US Centers for Disease Control and Prevention, albeit associated with severe infections (9). Notably most reports use genomic data from >5 y ago and there is a need to reappraise prevalence. The reported multi-country increase in group A streptococcal infections (1) since the lifting of pandemic restrictions underlines the importance of enhanced global surveillance for lineages with potentially enhanced fitness such as M1_UK_.

**Figure 1.**
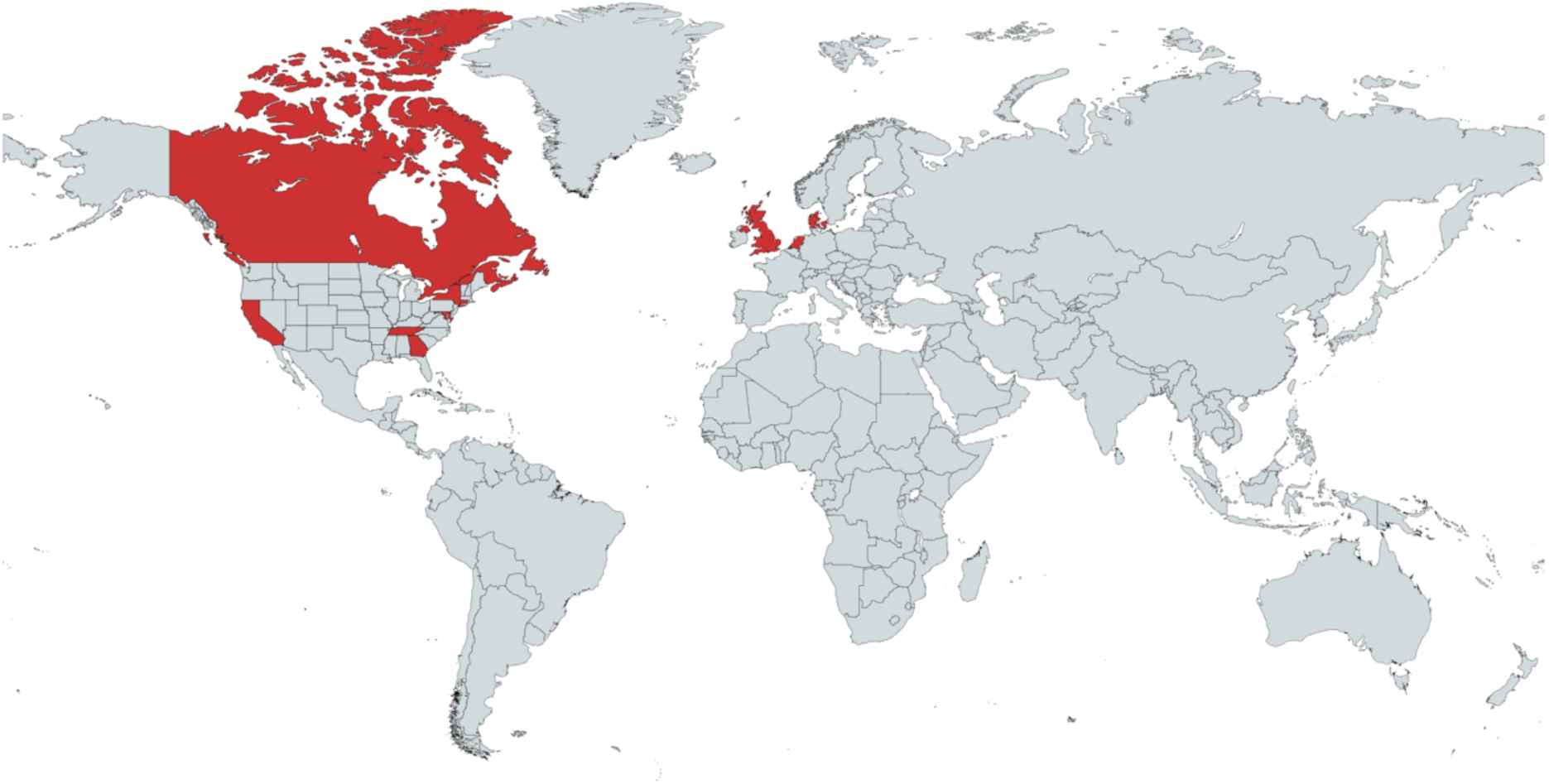
Countries or states (USA) with reported M1_UK_. Created with mapchart.net.

**Figure 2.**
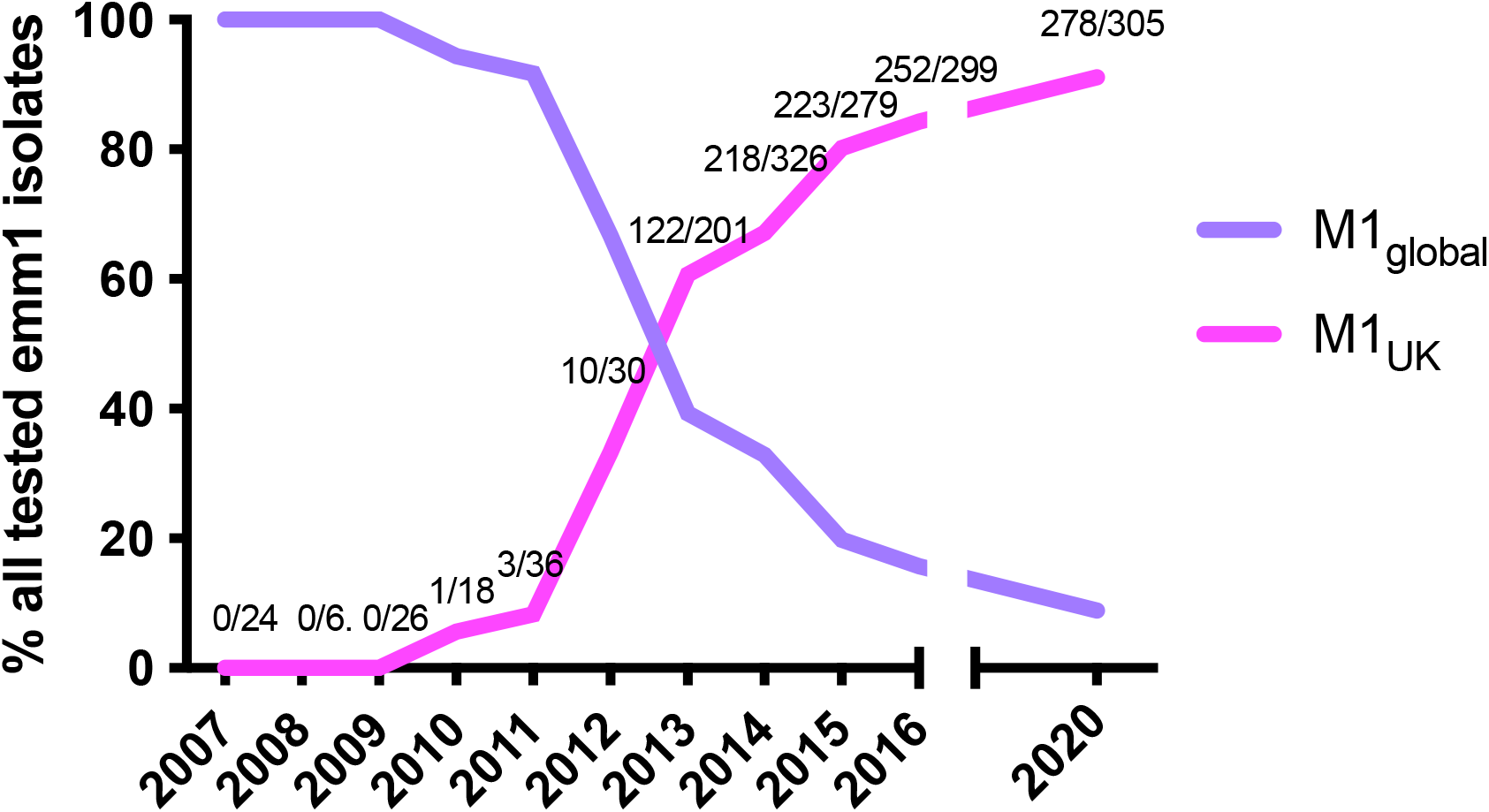
Proportion of *emm*1 isolates in England that belong to M1_UK_ using all available *emm*1 strains that were genome sequenced (2007-2016, data previously presented in Reference 4) and all available invasive isolates tested by allele-specific PCR (2020). Numbers on graph indicate number of isolates assigned as M1_UK_/total number tested in each year. Graph adapted and updated from Lynskey et al (Reference 4).

Genetic distinction of M1_UK_ and M1_global_ strains is possible using whole genome sequencing to detect the 27 SNPs that characterise the M1_UK_ lineage (4), however this is not available in all countries. We designed an allele-specific PCR (AS-PCR) assay to detect M1_UK_-specific SNPs in the following genes; *rofA, gldA* and *pstB*. Amplification targets were chosen to separate M1_UK_ and M1_global_ strains, but also to identify strains from less common ‘intermediate’ sublineages that include only 13 or 23 of the 27SNPs that characterise M1_UK_. (4). PCR conditions were optimised for each pair of amplicons using control strains from each lineage. (Table, supplementary figure 1).

**Table:**
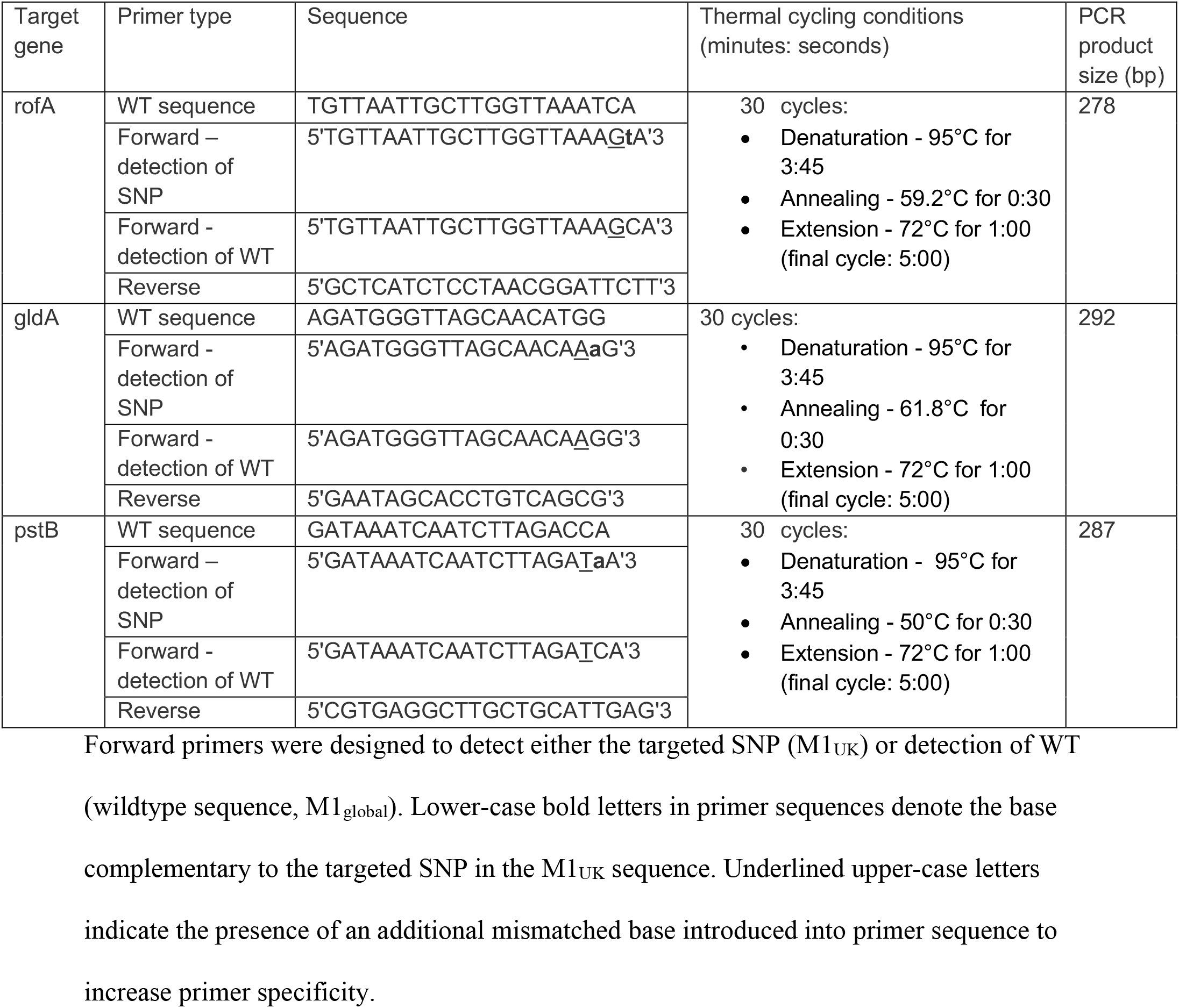
Allele specific PCR primers and conditions to differentiate M1_global_ and M1_UK_ S. pyogenes

To evaluate the allele-specific PCR, we tested if the *rofA* and *pstB* primers could correctly identify the lineage of 27 genome-sequenced *emm*1 *S. pyogenes* strains from 2017-2018 from the Imperial College bioresource of non-invasive *S. pyogenes* isolates. The isolates tested were artificially enriched for M1_global_ isolates to ensure the assay could distinguish the two lineages; 8/27 isolates were M1_global_, and 19/27 were M1_UK_. PCR amplification of *rofA, and pstB* alleles using DNA from these isolates assigned 19/19 M1_UK_ strains and 8/8 M1_global_ strains to the correct lineages. (Supplementary Table 1).

To validate the ability of AS-PCR to identify *emm*1 isolates from not only M1_global_ and M1_UK_, but also intermediate sublineages previously identified (4), a further 16 strains from 2014-2015 were tested, that comprised 4 strains each of M1_global_, M1_13snps_, M1_23snps_, and M1_UK_. SNPs were correctly detected in *rofA* in all strains belonging to M1_13snps_, M1_23snps_, and M1_UK_. SNPs were also correctly detected in *gldA* in all strains belonging to M1_23snps_ and M1_UK_, but not M1_global_, or M1_13snps_. Finally, SNPs in *pstB* were only identified in M1_UK_. (Supplementary Table 2).

In England, all isolates from invasive infection are submitted to the reference laboratory for *emm* typing. *Emm*1 isolates routinely represent the dominant genotype among invasive sterile site isolates, representing 20-30% of invasive infections. In 2020, a year where common respiratory infections including *S. pyogenes* were disturbed by COVID-19-related public health interventions, 305 invasive *emm*1 *S. pyogenes* isolates were available for study. AS-PCR identified SNPs in *rofA, gldA* and *pstB* in 278/305 (91.1%) of isolates that were therefore assigned as M1_UK_; while no SNPs were detected in the remaining 27 isolates, that were assigned as M1_global_. No intermediate lineage *emm*1 strains were identified in 2020. The longevity of emergent *S. pyogenes* lineages in a population is hard to predict; while an emm89_emergent_ acapsular lineage has disseminated globally (10), an emergent *emm*3 SpeC-producing lineage ceased to be detectable within a few years of first detection (11). Taken together with previously reported data combined from all available previously-sequenced *emm*1 isolates (Figure 1), the M1_UK_ lineage has continued to expand since 2016-end 2020 among invasive isolates in England.

The recent reports of increased invasive group A streptococcal activity in several countries (1) underline the importance of ongoing surveillance for novel lineages, given potential impact on public health. The AS-PCR reported herein provides a readily available method to detect M1_UK_; it is straightforward to undertake and, for screening purposes only, can be simplified to use only RofA primers, to rule out the presence of sublineages associated with M1_UK_, or M1_UK_. AS-PCR does not however replace genome sequencing as a gold standard for surveillance of highly pathogenic bacteria, albeit that sequencing is not widely available and can be expensive.

At the current time, *emm*1 strains account for over 50% of invasive infections in children in England (12). The data presented here confirm the M1_UK_ lineage to be dominant in England, with ongoing expansion to end of 2020, while contact tracing data from 2018 demonstrate a high frequency of secondary acquisition of M1_UK_ in school outbreak settings (13). Given the recognized association between *emm*1 *S. pyogenes* and outcome in invasive infection (14) enhanced surveillance for this sublineage in particular is warranted.

## Supporting information

Supplementary Figure and four tables

## Acknowledgements

The authors acknowledge the NIHR Imperial Biomedical Research Centre and NIHR Health Protection Research Unit in Healthcare-associated Infection (HPRU) and Antimicrobial Resistance at Imperial College London. The support of scientific and biomedical staff from the NIHR BRC Colebrook laboratory, NIHR HPRU, NIHR BRC Imperial Genomics Facility, and North-west London Pathology/Imperial College Healthcare Trust Diagnostic laboratory who contribute to the collection, biobanking, typing, and sequencing of bacterial isolates is also gratefully acknowledged.

