## Supplementary Figure and four tables for "Ongoing emergence of M1_UK_ lineage among invasive group A streptococcus isolates in 2020 and use of allele-specific PCR"

Xiangyun Zhi et al.

**Contents**

**Supplementary Figure** Gel electrophoresis demonstrating allele-specific PCR to distinguish $M1_{global}$ and $M1_{UK}$

**Supplementary Table 1** Evaluation of rofA and pstB allele-specific primers in lineage assignment, using genome-sequenced *emm*1 strains from 2017-2018

**Supplementary Table 2.** Validation of rofA, pstB, and gldA allele-specific primers in lineage assignment, using genome-sequenced *emm*1 strains including intermediate sublineages.

**Supplementary Table 3.** Use of AS-PCR to test 305 invasive *emm*1 *S. pyogenes* isolates of unknown lineage assignment

**Supplementary Table 4.** Genome-sequenced strains used to validate AS-PCR to detect all sublineages

**Supplementary Figure 1** Gel electrophoresis demonstrating allele-specific PCR to distinguish $M1_{global}$ and $M1_{UK}$

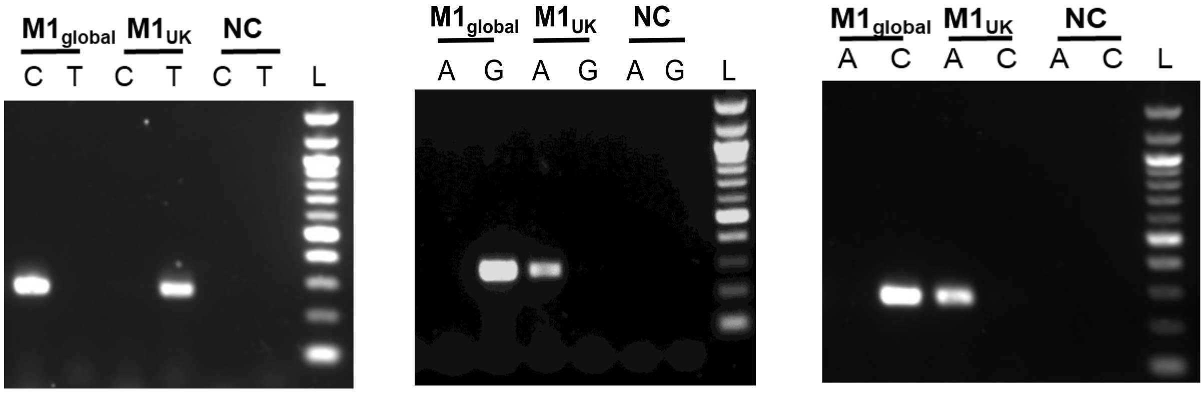


A. RofA primers B.gldA primers C. pstB primers

PCR products following amplification of DNA from $M1_{global}$ and $M1_{UK}$ strains of *S. pyogenes* are shown for (A) RofA SNP primers (C or T denotes use of either *rofA* C or T forward primers); (B) GldA SNP primers (A or G denotes use of either *gldA* A or G forward primers); (C) PstB SNP primers (A or C denotes use of either *pstB* A or C forward primers. DNA was from genome-sequenced control strains BHS0151 ($M1_{global}$) and BHS581 ($M1_{UK}$), listed in Reference 3; genome sequences are in the European Nucleotide Archive under accession references ERS1020136 and ERS1020603 respectively. Abbreviations, NC = negative control; L = 100bp DNA ladder.

.

**Supplementary Table 1.** Evaluation of rofA and pstB allele-specific primers in lineage assignment, using genome-sequenced *emm*1 strains from 2017-2018

| Year | WGS lineage assignment | Total no. of strains | Number of strains yielding PCR product with allele-specific primer pairs | | | |
|---|---|---|---|---|---|---|
| | | | RofA_$M1_{global}$ | RofA_$M1_{UK}$ | PstB_$M1_{global}$ | PstB_$M1_{UK}$ |
| 2017 | $M1_{global}$ | 3 | 3 | 0 | 3 | 0 |
| | $M1_{UK}$ | 6 | 0 | 6 | 0 | 6 |
| 2018 | $M1_{global}$ | 5 | 5 | 0 | 5 | 0 |
| | $M1_{UK}$ | 13 | 0 | 13 | 0 | 13 |

AS-PCR results performed on extracted DNA were consistent with genome sequencing (WGS) of DNA from 27 genome sequenced *emm*1 non-invasive *S. pyogenes* isolates (n=9 from 2017, and n=18 from 2018, enriched for M1global strains).
Sequences for *emm*1 test strains submitted to European nucleotide archive, accession records for sequenced strains in process.

S**upplementary Table 2.** Validation of rofA, pstB, and gldA allele-specific primers in lineage assignment, using genome-sequenced *emm*1 strains including intermediate sublineages.

| WGS lineage assignment | Total no. of strains | Number of strains yielding PCR product with allele-specific primer pairs | | | | | |
|---|---|---|---|---|---|---|---|
| | | RofA $M1_{global}$ | RofA $M1_{UK}$ | GldA $M1_{global}$ | GldA $M1_{UK}$ | PstB $M1_{global}$ | PstB $M1_{UK}$ |
| $M1_{global}$ | 4 | 4 | 0 | 4 | 0 | 4 | 0 |
| $M1_{13snp}$ | 4 | 0 | 4 | 4 | 0 | 4 | 0 |
| $M1_{23snp}$ | 4 | 0 | 4 | 0 | 4 | 4 | 0 |
| $M1_{UK}$ | 4 | 0 | 4 | 0 | 4 | 0 | 4 |

16 whole genome-sequenced *S. pyogenes emm1* strains from 2013 and 2014 (Accession numbers, Supplementary Table 4) were analysed using RofA, GldA and PstB AS-PCR. Strains were selected for the presence of 0/27snps; 13/27; 23/27snps; or 27/27 SNPs (4 strains from each). AS-PCR results were consistent with sequencing.

**Supplementary Table 3.** Use of AS-PCR to test 305 invasive *emm*1 *S. pyogenes* isolates of unknown lineage assignment submitted in 2020

| | RofA $M1_{global}$ | RofA $M1_{UK}$ | GldA $M1_{global}$ | GldA $M1_{UK}$ | PstB $M1_{global}$ | PstB $M1_{UK}$ | Inferred lineage |
|---|---|---|---|---|---|---|---|
| No. strains yielding PCR product with AS-primers | 27 | 0 | 27 | 0 | 27 | 0 | $M1_{global}$ n=27 |
| | 0 | 278 | 0 | 278 | 0 | 278 | $M1_{UK}$ n=278 |

All invasive emm1 strains (n=305) submitted to the reference laboratory for *emm* typing and identified as *emm*1 were evaluated by AS-PCR as shown above. 278 isolates yielded PCR products consistent with $M1_{UK}$. No intermediate isolates were identified.

**Supplementary Table 4. Accession numbers of genome-sequenced strains used to validate AS-PCR to detect all sublineages**

| M1-lineage | Year | ERR number¶ | Sample ID¶ |
|---|---|---|---|
| 13SNPs | 2013 | ERS1594714 | PHEGAS005 |
| 13SNPs | 2013 | ERS1594852 | PHEGAS127 |
| 13SNPs | 2015 | ERR1733723 | GASEMM2799 |
| 13SNPs | 2015 | ERR1734520 | GASEMM2970 |
| 23SNPs | 2013 | ERS1594734 | PHEGAS025 |
| 23SNPs | 2013 | ERS1594744 | PHEGAS035 |
| 23SNPs | 2013 | ERS1594757 | PHEGAS048 |
| 23SNPs | 2013 | ERS1594864 | PHEGAS137 |
| M1global | 2013 | ERS1594798 | PHEGAS168 |
| M1global | 2013 | ERS1594822 | PHEGAS097 |
| M1global | 2014 | ERR1732733 | GASEMM1027 |
| M1global | 2015 | ERR1734897 | GASEMM2755 |
| M1uk | 2013 | ERS1594722 | PHEGAS013 |
| M1uk | 2014 | ERR1733140 | GASEMM0629 |
| M1uk | 2015 | ERR1733678 | GASEMM3027 |
| M1uk | 2016 | ERS1594947 | PHEGAS285 |

¶Sequences in European Nucleotide Archive; previously listed in Reference 4.
